# NHA1 is a cation/proton antiporter essential for the water-conserving functions of the rectal complex in *Tribolium castaneum*

**DOI:** 10.1101/2022.09.29.510179

**Authors:** Muhammad Tayyib Naseem, Robin Beaven, Takashi Koyama, Sehrish Naz, Mooney Su, David P. Leader, Dan Klærke, Kirstine Calloe, Barry Denholm, Kenneth Veland Halberg

## Abstract

More than half of all extant metazoan species on earth are insects. The evolutionary success of insects is intrinsically linked with their ability to osmoregulate, suggesting that they have evolved unique physiological mechanisms to maintain water balance. In beetles (Coleoptera)—the largest group of insects—a specialized rectal (‘cryptonephridial’) complex has evolved that recovers water from the rectum destined for excretion and recycles it back to the body. However, the molecular mechanisms underpinning the remarkable water-conserving functions of this system are unknown. Here, we introduce a transcriptomic resource, BeetleAtlas.org, for red flour beetle *Tribolium castaneum*, and demonstrate its utility by identifying a cation/H^+^ antiporter (NHA1) that is enriched and functionally significant in the *Tribolium* rectal complex. NHA1 localizes exclusively to a specialized cell type, the leptophragmata, in the distal region of the Malpighian tubules associated with the rectal complex. Computational modelling and electrophysiological characterization in *Xenopus oocytes* show that NHA1 acts as an electroneutral K^+^/H^+^ antiporter. Furthermore, genetic silencing of *Nha1* dramatically increases excretory water loss and reduces organismal survival during desiccation stress, implying that NHA1 activity is essential for maintaining systemic water balance. Finally, we show that Tiptop, a conserved transcription factor, regulates NHA1 expression in leptophragmata and controls leptophragmata maturation, illuminating the developmental mechanism that establishes the novel functions of this cell. Together, our work provides the first insights into the molecular architecture underpinning the function of one most powerful water-conserving mechanisms in nature, the beetle rectal complex.

**Significance Statement:** Beetles are the largest group of insects, inhabiting a wide range of habitats on earth. Unique adaptations in overcoming water stress is critical to their success, yet the mechanisms underpinning this ability are unknown. Using genetics, electrophysiology, imaging and behavioral studies we show that a cation/H^+^ (NHA1) transporter is exclusively localized to specialized cell type, the leptophragmata, in the Malpighian tubules associated with the rectal complex. Ion transport functions of NHA1 in leptophragmata underpin the movement of water from the rectum, from where it would be destined for excretion, to the Malpighian tubule and then recycled back to the body. This water recovery capability of rectal complex is essential for maintaining systemic water balance in beetles. This work provides the first insight into to the molecular architecture of one of most powerful water-conservation mechanisms in biology, and provides an important clue to the ecological and evolutionary success of the beetles.

## Introduction

Insects are (in terms of species) the most diverse animal group on the planet, occupying the widest possible range of habitats on earth (1). However, insects—being a predominantly terrestrial group—are faced with major problems in maintaining ion and water balance as their small size and large surface-to-volume ratio makes them highly sensitive to osmotic disturbances. As such, the evolutionary success of insect is intrinsically linked with their ability to defend against harmful changes in their water contents in a wide range of environments. In insects, the renal (Malpighian) tubules (MTs) and the hindgut are the principle organs responsible for regulating body fluid composition (2). Whereas the MTs secrete excess ions and water by producing a primary urine that is drained into the alimentary canal (3), the hindgut selectively reabsorbs solutes and water in proportion to the needs of the animal (4). The hindgut (in particular the rectum) is thus the major site of water conservation in insects as it provides vital feedback control of the final composition and volume of the excretory products. However, surprisingly little is known about the mechanisms that mediate the selective absorption of water and solutes by the rectum in different external conditions.

The physiological importance of the rectum in maintaining overall water balance in the insect is particularly evident in species colonizing osmotically hostile environments. For example, in the mealworm *Tenebrio molitor*—a species that is capable of completing its entire life cycles without access to environmental water—a specialized rectal (‘cryptonephridial’) complex has evolved that enables recovery of almost all water from the rectum (5, 6). The anatomical arrangement of this complex is defined by the distal ends of the MTs (perirectal tubules, PTs) being closely applied to the rectal wall with the entire structure enclosed beneath an impermeable perinephric membrane (6, 7). The water-conserving properties of the rectal complex rely on the active KCl transport by the PTs to generate a fluid of sufficiently high osmotic pressure to facilitate osmotically-driven water removal from the feces (8, 9). In effect, the system establishes a standing gradient along the anterior-posterior axis of the complex to maximize fluid absorption in a manner analogous to the countercurrent arrangement of the vertebrate nephron in the renal medulla (10). In this way, *Tenebrio* (and its relatives) is able to extract and recycle almost all water from the rectal lumen to produce powder-dry excreta (7, 11). Remarkably, the system can even be used as a novel physiological mechanism for water uptake by enabling absorption of water vapor directly from the moist air (5–7, 11, 12). It has been suggested that the accumulation of KCl in the PTs is mediated by a small population of secondary cells (SCs) known as ‘leptophragmata’, as they are the only cells that interrupt the perinephric membrane to enable hemolymph-to-tubule movement of KCl (6, 7). However, in spite of having been intensely studied for almost a century, the cellular and molecular architecture underpinning the water-extracting functions of this extraordinary organ remains entirely unknown.

Cataloging the relative strength and specificity of gene expression across different tissues and life stages of an organism can provide valuable insights into most biological functions. Transcriptomic atlases have therefore become powerful tools in the functional genomics arsenal by enabling the annotation of physiological mechanisms and developmental processes on a gene-by-gene basis (13–17). However, despite their obvious benefits to their respective communities, the number of such transcriptomic atlases is limited. In the field of insect functional genomics, the red flour beetle *Tribolium castaneum* (a closely related species of *Tenebrio)* has emerged as a powerful model system because of its rapidly expanding transgenic toolkit (18, 19) and its amenability to large-scale mutagenic studies (20, 21). The construction of an authoritative expression atlas for *Tribolium* would thus not only complement with existing post-genomic resources, but also help broaden the scope for functional analysis of one of the most economically (contain many devastating crop pests) and ecologically (largest group of insects) important animal groups on Earth, the beetles.

Here we report the development of BeetleAtlas, a transcriptomic atlas covering distinct ontogenetic and tissue-specific expression profiles of *Tribolium,* and demonstrate the utility of this resource by identifying a cation/H^+^ antiporter (NHA1) that is essential to the water-extracting properties of the *Tribolium* rectal complex. Using a combination of bioinformatics, genetics, imaging, electrophysiology, and organ assays we show that NHA1 localizes exclusively to the specialized leptophragmata cells in the PTs where it acts as an electroneutral cation/H^+^ antiporter. Genetic depletion of *Nha1* dramatically increases excretory water loss and impairs whole-animal survival during desiccation stress, suggesting that NHA1 activity is essential for maintaining systemic water balance. Finally, we show that NHA1 expression and leptophragmata maturation is regulated by a transcription factor called Tiptop, which is part of a conserved gene regulatory network that is central to the function of the rectal complex in tenebrionid beetles. Taken together, our work provides the first insights into the molecular architecture underlying the function of one most powerful water-extracting mechanisms in biology, the tenebrionid rectal complex.

## Results

### BeetleAtlas: A transcriptional atlas of *Tribolium* tissues and life stages

A large fraction of any animal genome is differentially expressed in distinct cell types, tissues and life stages. Therefore, it is vital to consider spatio-temporal changes in gene expression in order to understand cell functions. To gain a comprehensive view of the transcriptional landscapes of individual tissues and life stages of *Tribolium,* we have created BeetleAtlas (BeetleAtlas.org). This is a transcriptional atlas of gene expression based on RNA sequencing (RNAseq) that covers several embryonic stages as well as major larval and adult tissues. Specifically, it catalogs transcriptomic data sets covering 11 distinct adult tissues (head, brain, anterior midgut, posterior midgut, hindgut, Malpighian tubules, rectal complex, fat body, female /male gonads and carcass) dissected from 7-day-old San Bernardino (SB) adult beetles; nine larval tissues (head, brain, anterior midgut, posterior midgut, hindgut, Malpighian tubules, rectal complex, fat body and carcass) dissected from L6 larvae; and 4 distinct (0-1, 1-24, 24-36 and 36-72 hours post egg lay) stages of embryonic development (Fig. 1*A*). Each sample was prepared according to standardized protocol and sequenced on the same sequencing platform in a minimum of three biological replicates and compared to matched whole-animal samples. Together, BeetleAtlas thus allows systematic comparisons of gene expression across all major tissues and ontogenetic stages of the more than 16,500 genes (with over 18,500 transcripts) encoded by the *Tribolium* genome (22).

**Fig 1.**
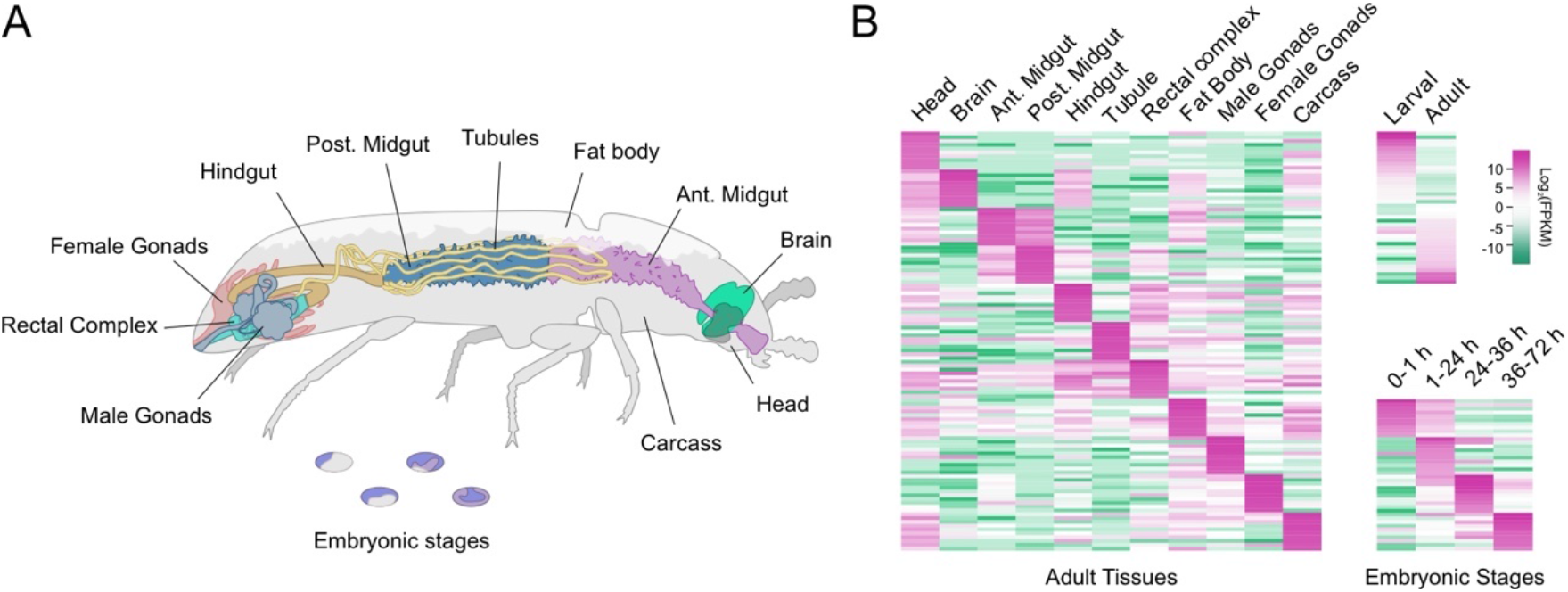
BeetleAtlas is an authoritative transcriptomic atlas of gene expression. (*A*) Adult *Tribolium* anatomy highlighting the tissues selected for micro-dissection and bulk RNA-sequencing. (*B*) Clustered heat maps of gene expression (log_2_ transformed values of fragments per kilobase of transcript per million, FPKM) across the tissue-specific and ontogenetic samples demonstrating distinct transcriptional signatures.

An implicit requirement of such a facility is the robustness of the data in the underlying database. To formally assess the quality of our data, we identified the most highly expressed genes in each sample and generated a clustered heat map to visualize the divergence of transcriptomes between the different ontogenetic stages and larval and adult tissues (Fig. 1*B*). These data show a clear discrimination of the transcriptional signatures, not only between different life stages, but also between physically adjacent tissues within the same stage, such as the larval midgut and tubules or the adult rectal complex and gonads. Furthermore, we performed manual searches for genes that show a strong enrichment in a particular tissue and performed independent validation of gene expression by qRT-PCR. These results further confirm and validate the spatial expression profiles reported by BeetleAtlas across the selected genes (*SI Appendix*, Table 1). BeetleAtlas is thus a powerful tool for the *Tribolium* and insect communities, providing a critical first step by mapping gene activity to particular stages or tissues and by offering valuable information on the function of each gene product by integrating complementary large-scale RNA interference (RNAi) screening data (20).

### BeetleAtlas identifies candidate genes involved in rectal complex function

To demonstrate the utility of BeetleAtlas in providing transcriptional insights into tissue function, we aimed to characterize the molecular mechanisms underpinning the function of the rectal (‘buried kidney’) complex (6). Using the ‘Gene’ look-up section of BeetleAtlas, we performed manual searches of all genes identified as core components of the insect ‘epitheliome’ (23) and looked for transcript enrichment of the *Tribolium* orthologues relative to that of the whole animal. In addition, we used the ‘Tissue’ search function to perform an unbiased search for candidate transporter genes that are most highly enriched in the rectal complex. Adopting parallel hypothesis-driven (candidate genes) and hypothesis-free (gene enrichment) approaches, we identified more than 30 putative genes that are expressed at very high levels in this multiorgan system compared to the organismal average, and further revealed that most of these genes belong to a core set that is co-expressed of other insect transport epithelia (23, 24). From this candidate list, *TC013096*, a gene predicted to encode a cation/H^+^-antiporter (*Nha1*), shows the highest enrichment in the rectal complex (Fig. 2*A*). Indeed, mapping the tissue-specific expression pattern of *Nha1* indicates that this gene is predominantly expressed in the rectal complex of both larval and adult *Tribolium* (Fig. *2B-D*); an expression profile that we validated by qRT-PCR (*SI Appendix*, Table 1). Together, the transcriptional signature of *Nha1* suggests that this transporter is likely to play a key role in the function of the *Tribolium* rectal complex.

**Fig 2.**
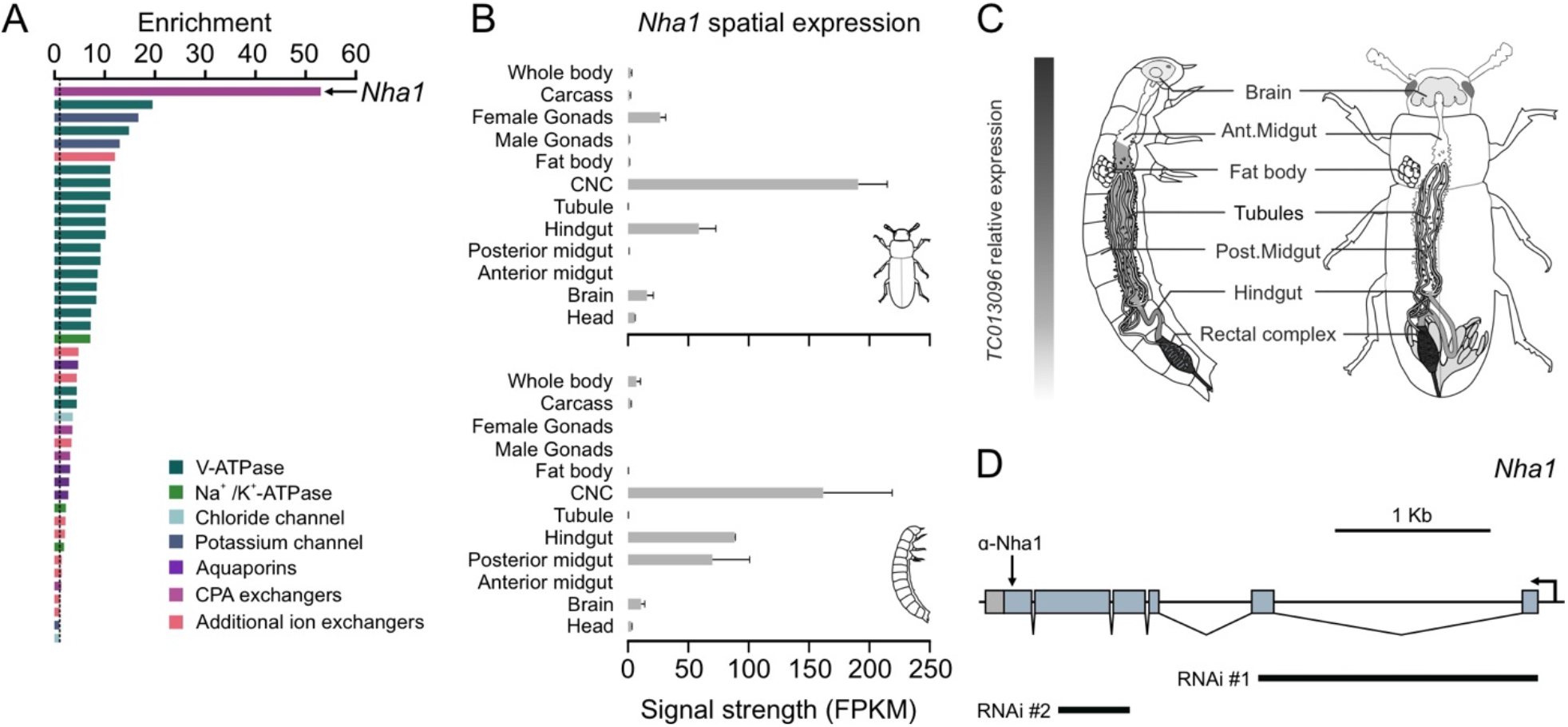
Mining BeetleAtlas to identify novel molecules involved in rectal complex function. (*A*) Fold increase (enrichment) in gene expression of membrane channels and transporters in the rectal complex relative to the whole animal (dashed line) according to BeetleAtlas. A gene encoding a putative Nha1-like protein is most highly enriched in the CNC. (*B*) Spatial expression analysis of *Nha1* shows that it is almost exclusively expressed in the hindgut and CNC of both larval and adult *Tribolium. (C*) Heat map of *Nha1* expression superimposed on larval and adult anatomy. (*D*) Predicted exon map of *Nha1* marking the epitope targeted for antibody generation and regions selected for dsRNA synthesis. Of the two fragments tested, fragment #2 produced the strongest knockdown and was therefore used in the remaining part of the study.

### NHA1 localizes to a specialized cell type in the perirectal tubules of the rectal complex

To gain insight into the physiological roles of NHA1 we next examined the structure and spatial organization of the *Tribolium* rectal complex using scanning electron microscopy. Like the rectal complex from the closely related mealworm *Tenebrio* (6, 7, 25), the *Tribolium* rectal complex is organized according to the ‘cryptonephridial’ condition. This condition is characterized by the distal ends of the MTs being closely applied to the rectal epithelium in a sinuous pattern and enclosed within a compartment, the perinephric space that is separated from the body cavity by an impermeable perinephric membrane (Fig. 3*A-C*). Notably, the distal regions of the MTs—the so-called perinephric tubules (PTs)—are conspicuously different from the ‘free’ part of the tubules as they are characterized by small dilations over which the perinephric membrane is extremely thin and in direct contact with the dilations (‘boursourflors’) under which specialized small-nucleated cells known as leptophragmata are found (Fig. *3B-B*) The anatomical position of these cells is intriguing as previous studies in *Tenebrio* had demonstrated that the function of the rectal complex is based on the accumulation of high concentrations of KCl in the PT lumen mediated by active hemolymph-to-tubule transport to facilitate osmotically driven water removal from the feces (6, 7, 9). Such a model is supported by the high expression of genes encoding the different subunits of the plasma membrane Vacuolar H^+^ ATPase (V-ATPase) (Fig. *2A*; *SI Appendix* Fig. S1*C*)—a plasma membrane transporter critical to energizing insect epithelia (26)—and by the exclusive localization of the protein to the apical brush border of the PTs in the *Tribolium* rectal complex (Fig. 3*G*). As the perinephric membrane is highly impermeable except for the ‘blister-like’ windows under which leptophragmata sit (27), the anatomical data strongly suggest that these small cells are the only sites of exchange. Consistent with this notion, backscattered electron detection (BSE)—a method that carries information on differences in atomic number (*Z*) of the sample—on rectal complexes preincubated in silver nitrate (27), reveals that silver staining is exclusively found at sites corresponding to the anatomical position of the leptophragmata (Fig. *3A-F*), implying that hemolymph-to-tubule transport is exclusively mediated by these cells. Strikingly, NHA1 protein was shown to localize entirely to a small-nucleated cell type resembling the leptophragmata of the PTs (Fig. *3H*) using an NHA1-specific antibody. The specificity of the antibody was verified by lack of immunoreactivity in *Nha1* depleted animals (*SI Appendix*, Fig. *1A-B*). Indeed, the expression of NHA1 in leptophragmata was confirmed by double staining with Tiptop (Tio) (Fig. *3I*), a transcription factor that marks leptophragmata and is involved in the differentiation of secondary cells (SCs) in *Tribolium* and other insects (28–30). This finding suggests that the leptophragmata and the SCs of the free tubules are related. Taken together, our data indicate that the anatomical features of the *Tribolium* rectal complex are largely identical to those described previously for *Tenebrio* (6, 7), and that NHA1 is exclusively localized to the specialized leptophragmata cells in the PTs of the rectal complex.

**Fig 3.**
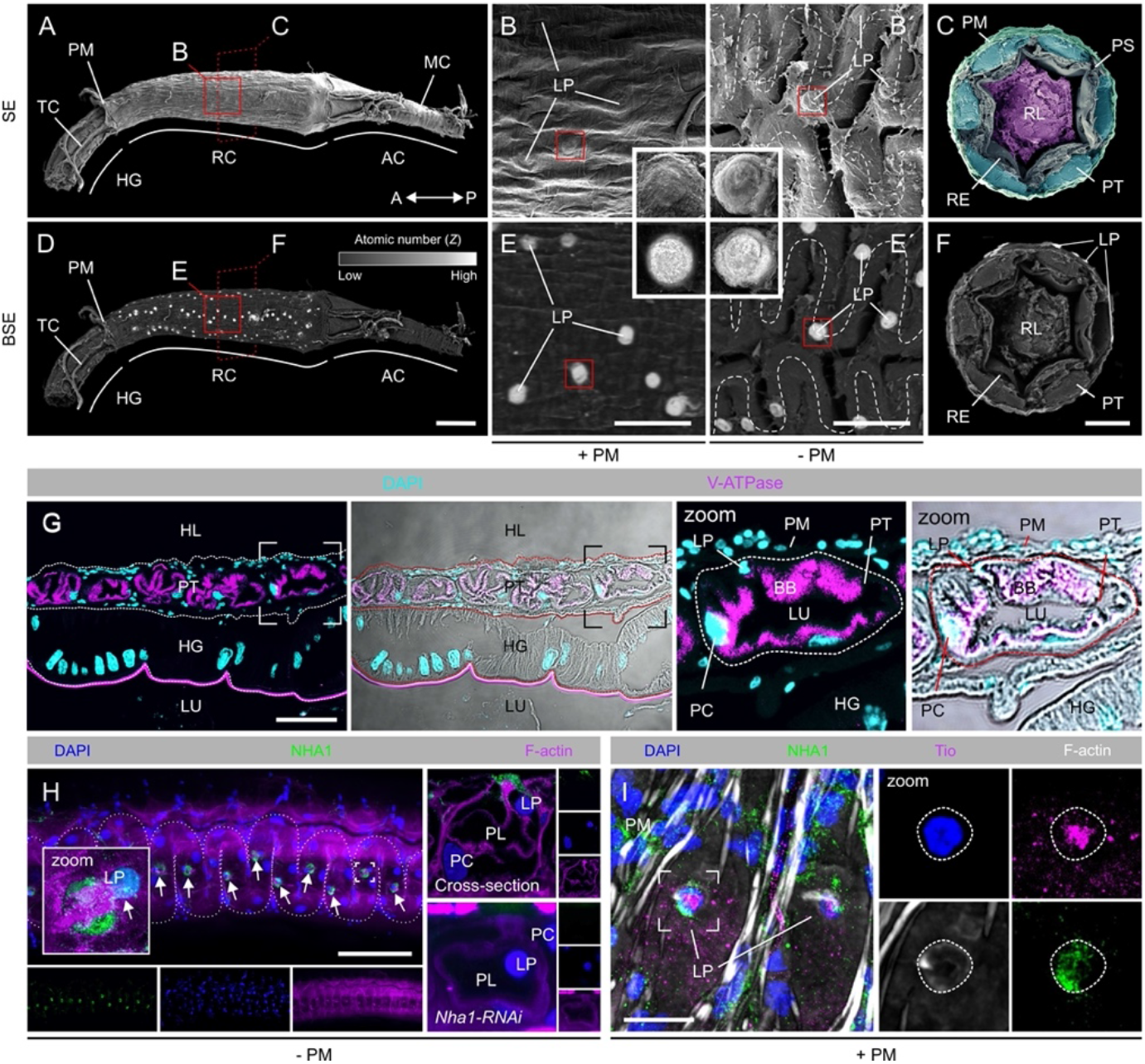
NHA1 localizes to specialized leptophragmata cells in the PTs of the rectal complex. (*A-C*) Scanning electron microscopy (SEM) images showing the gross morphology of the rectal complex. A subpopulation of cells is arranged along the PTs as a series of dilations that extend radially towards the PM suggesting that they are leptophragmata. (*D-F*) Back-scattered electron detection reveal that these dilations contain elements of high atomic number (*Z*) following AgNO3 application confirming that these dilations are LP. (*G*)Maximum projection confocal microscopy images of paraffin-sections of the rectal complex demonstrate that the V-ATPase localizes almost exclusively to the apical brush-border of PCs in the PTs. (Scale bar, 40 μm). Note the cuticle of the rectal epithelium shows strong autofluorescence. (*H*) Maximum projection confocal microscopy images demonstrate that NHA1 is expressed in this subpopulation of LP cells along the PTs (small arrows). Subcellular localization of NHA1 demonstrates exclusive expression to the small-nucleated LP cells (cross-section of a PT). Rectal complex dissected from animals injected with dsRNA targeting the *Nha1* gene (*Nha1-RNAi*)shows a dramatic reduction in immunoreactivity confirming the specificity of the antibody. (Scale bar, 100 μm.) (*I*) LP identity is further confirmed by co-expression of the Tiptop (Tio) transcription factor, which is a marker of the LPs (as well as secondary cells in the “free” part of the tubule (28)). AC, anal canal; HG, hindgut; LP, leptophragmata; MC, muscle cell; PC, principal cell; PM, perinephric membrane; PS, perinephric space; PT, perinephric tubule; RC, rectal complex; RL, rectal lumen; RE, rectal epithelium; TC, trachea. BB, brush-border; HG, hindgut; HL, hemolymph; PC, principal cell; PM, perinephric membrane; PT, perinephric tubule; LP, leptophragmata; LU, lumen.

### NHA1 acts as an electroneutral cation/H^+^ antiporter

The NHA transporters belong to the Cation/Proton Antiporter-2 (CPA2) subfamily of proteins, which are known to participate in a broad range of transport mechanisms across all kingdoms of life (31). Classically, the NHAs are secondary active transporters that exchange sodium ions against protons, yet several CPA2 members have been shown to move different types of substrates (32, 33), suggesting that their molecular function cannot be inferred from structural similarity. To gain insight into the role of NHA1 in rectal complex physiology we first performed sequence analysis and computational modelling of the protein. Surprisingly, we found only one *Nha* gene (*Nha1*) encoded by the *Tribolium* genome, which contrasts with other insect genomes in which two paralogs (*Nha1* and *Nha2*) are present (31, 32, 34); a BLASTp search against *Drosophila* proteins reveals that *Tribolium* NHA1 is more closely related to *Drosophila* NHA1 (53% sequence identity) than to NHA2 (35% sequence identity). Next, we performed three-dimensional structure predictions using I-TASSER (35) followed by Ramachandran plot and molecular dynamics (MD) simulations (36) to identify the putative transmembrane domains and globular structure of the protein. Consistent with the structural hallmarks of CPA2 members, the structural architecture of *Tribolium* NHA1 is defined by 12 transmembrane helices (both the N- and C-terminal tails are located in the cytoplasm) that collectively form a negatively charged transport funnel containing the putative ion binding and translocation domains (Fig. 4*A-B*; *SI Appendix*, Fig. 2*A-C*). These findings led us to perform MD simulations in the presence of cation substrates (K^+^ or Na^+^) to help resolve the ion binding and cation selectivity of NHA1. After an initial phase of equilibrium, we observed that the structural stability of the protein was predicted to be higher (i.e. lower Radius of Gyration and Root Mean Square Deviation) in the presence of bound K^+^ relative to bound Na^+^ (Fig. 4*C-D*), implying that K^+^ binding is favored over Na^+^ by the *Tribolium* NHA1 protein.

**Fig 4.**
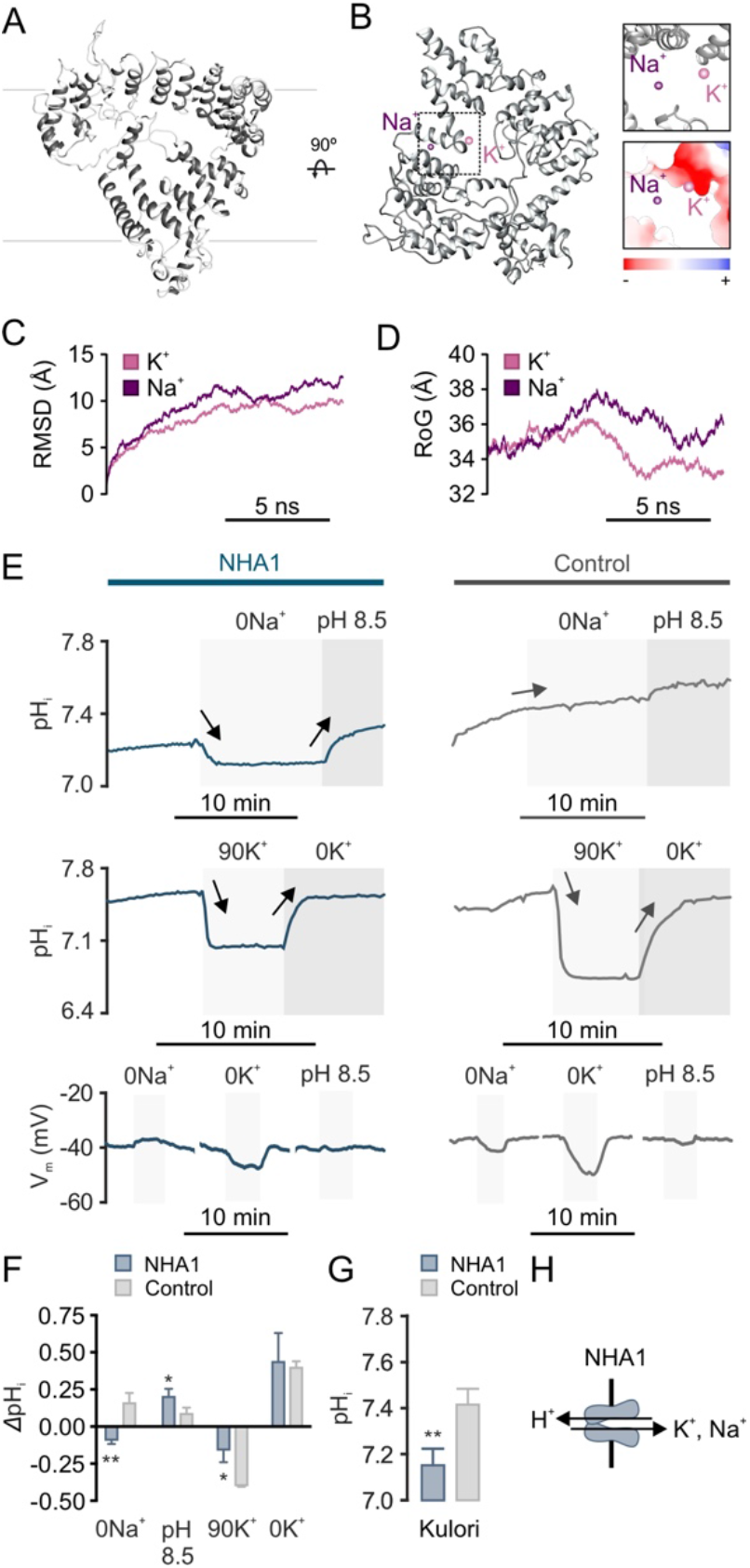
NHA1 acts as a K^+^(Na^+^)/H^+^ exchanger. (*A*) Predicted three-dimensional ribbon diagram of the NHA1 transporter embedded in the plasma membrane (*B*) Top view of the NHA1 protein highlighting K^+^ (pink) and Na^+^ (purple) binding; K^+^ is predicted to bind closer to negatively charged transport funnel compared to Na^+^(inserts). (*C*) Radius of gyration (RoG) plot of NHA1 compactness in the presence of K^+^ (pink) and Na^+^(purple) as a function of time. (*D*) Root mean square deviation (RMSD) plot of Nha1 backbone (C-α) atoms in the presence of K^+^ (pink) and Na^+^ (purple) as a function of time. (*E*) Representative traces of pHi and Vm of control (dark grey) or *Nha1* (blue) injected *Xenopus laevis* oocytes in response to changes in bath pH, [Na^+^] and [K^+^]. Significant changes in pHi are indicated by arrows. (*F*) Quantitative comparison of changes in pHi of oocytes in response to different challenges (Student’s *t*-test; n=5-10, * P<0.05, ** P<0.01). (*G*) Resting levels of pHi of oocytes from Nha1 (blue) and water-injected controls (dark grey) after incubation in Kulori’s solution pH 7.4 (Student’s *t*-test; n=5-10, ** P<0.01). (*H*) Schematic representation of ion transport mediated by NHA1.

We next sought to experimentally validate our MD simulations by expressing *Tribolium* NHA1 in *Xenopus* oocytes to allow electrophysiological characterization of the protein. Monitoring intracellular pH (pHi) in NHA1 oocytes while sequentially manipulating extracellular [Na^+^], [K^+^] and pH revealed that they rapidly responded to these changes, whereas similar responses were not observed in control oocytes injected with water (Fig. *4E-F*). These data suggest that *Tribolium* NHA1 is capable of recognizing both Na^+^and K^+^ as substrates. However, the significantly lower pHi of resting NHA1 oocytes relative to water-injected controls supports our *in silico* analysis that K^+^ is likely favored over Na^+^ (Fig. 4*G*). The membrane potential (Vm) of NHA1 oocytes was insensitive to extracellular pH implying that NHA1 transport is not electrogenic (Fig. 4*E*; *SI Appendix*, Fig. S3). Taken together, our results show that in *Tribolium* NHA1 is functionally coupled to the H^+^ gradient to mediate electroneutral exchange of K^+^ (Na^+^) for H^+^ (Fig. *4H*) in a manner similar to that observed for fungal KHA members (37–39).

### Internal water abundance modulates NHA1 activity in leptophragmata cells

The potent activation of the rectal complex in animals deprived of water (6) implies that the transport machinery of the complex is regulated in response to changes in hemolymph osmotic pressure. We therefore asked whether NHA1 activity is altered in animals exposed to conditions known to affect internal water abundance (28, 40). Quantifying NHA1 expression in the rectal complex revealed that both transcript and protein levels were consistently decreased in animals exposed to conditions that promote fluid retention (water, relative humidity, RH 90%), and significantly increased in beetles exposed to severe desiccation (RH 5%), compared to control animals (Fig. 5*A-B*). Furthermore, artificial activation (DH37 injection) and genetic deactivation (*Urn8* knockdown) of a potent diuretic pathway known to regulate organismal water levels (28) induced a robust increase and a significant decrease in *Nha1* expression, respectively (Fig. 5*C-D*). These observations suggest that NHA1 abundance is regulated in response to internal changes in hemolymph concentration as part of homeostatic mechanism that modulates the reabsorptive capacity of the rectal complex to maintain ion and water balance. Such a role is consistent with the observation that the concentration of the perirectal fluid in animals exposed to desiccation increases disproportionately relative to the hemolymph, presumably to allow more effective reabsorption of water from the rectal lumen (6).

**Fig 5.**
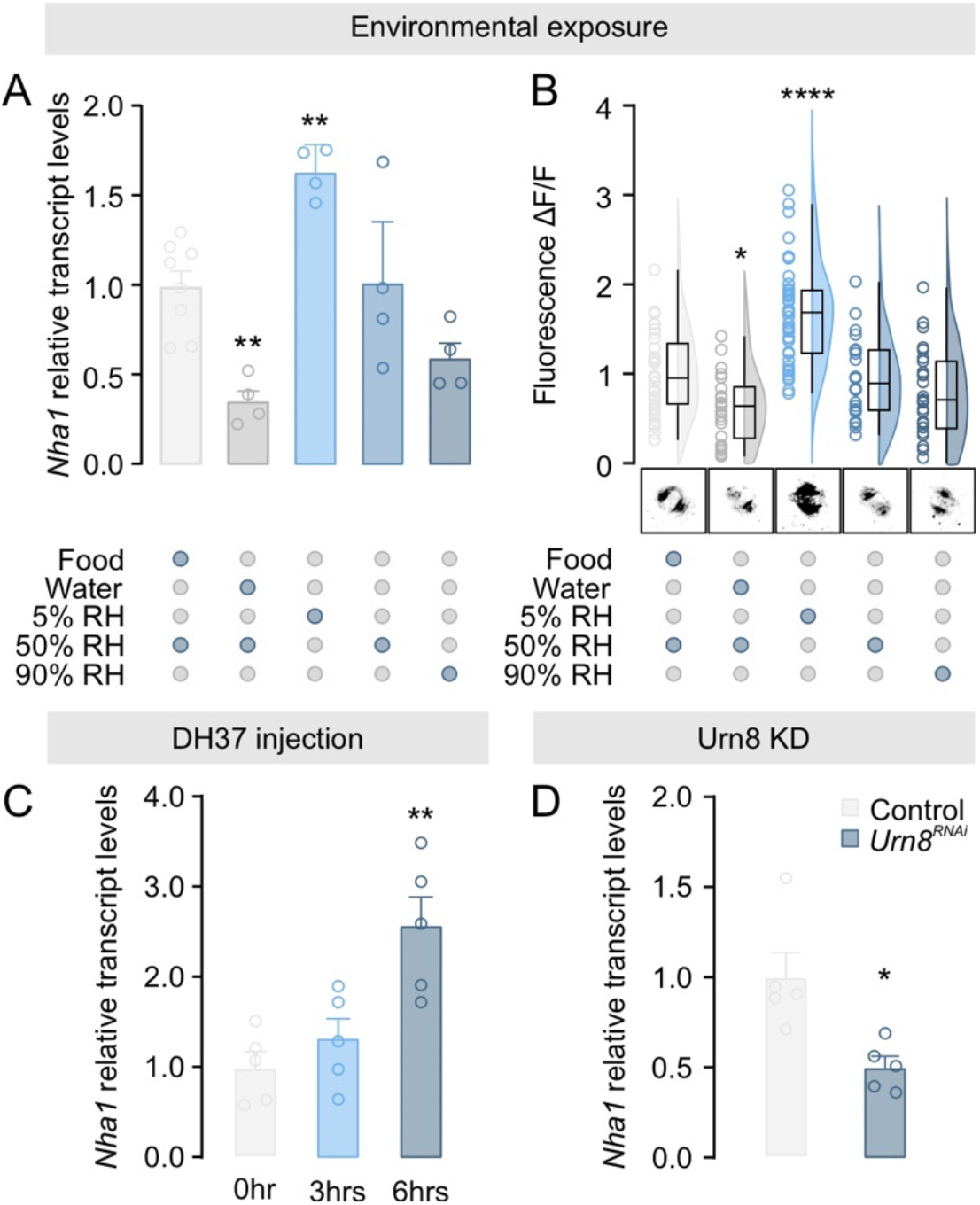
Environmental cues modulating rectal complex *Nha1* expression. (*A) Nha1* transcript levels in the rectal complex (*n* = 4-8) and (*B*) Raincloud plot of anti-NHA1 immunofluorescence (*n* = 23-45) in leptophragmata from animals exposed to different environmental conditions. Representative images of NHA1 immunofluorescence levels following exposure to each condition are shown below. Significant differences indicate pair-wise comparisons between control (food, RH 50%) and a given experimental group (one-way ANOVA, * *P*<0.05, **** *P*<0.0001). (*C*) *Nha1* transcript levels in the rectal complex (*n* = 5) from animals injected with DH37 (one-way ANOVA, ** *P*<0.01) or (*D*) from beetles injected with dsRNA targeting the *Urn8* gene (unpaired Student’s *t* test, * P<0.05).

### Silencing *Nha1* expression increases sensitivity to desiccation by inducing excessive water loss

We next sought to explore the functional significance of NHA1 in maintaining systemic water balance *in vivo* by selective downregulating *Nha1* expression using RNAi. Given that the rate of water loss is a crucial factor in determining the tolerance to desiccation in *Tribolium* and other insects (28, 41, 42), we hypothesize that *Nha1* depletion might lead to increased excretory fluid loss and thus impair the ability of *Tribolium* to survive dry conditions. Consistent with this hypothesis, *Nha1* silenced animals showed an increased sensitivity to desiccation relative to control injected animals, with a median survival of 3.3 days as compared to 5.8 days (Fig. 6*A*); RNAi efficacy was verified by qRT-PCR and immunocytochemistry showing >95% knockdown and a complete loss of detectable NHA1 expression (*SI Appendix*, Fig. S1*A-B*). This reduction in median survival is likely explained by an impaired ability to conserve water, since *Nha1* knockdown beetles consistently showed an increased rate of organismal water loss and a concomitant increase in hemolymph osmotic pressure (Fig. 6*B-C*). These effects were retained, albeit to a diminished extent, in animals exposed to high humidity implying that NHA1 function is essential to maintain systemic water balance across a wide range of conditions of fluid stress (*SI Appendix*, Fig. S4*A-C*). To test if the observed sensitivity to desiccation in *Nha1-*silenced beetles can be explained by increased excretory water loss, we examined the fecal output profiles of control and knockdown animals using an established *in vivo* excretion assay based on a dye-laced food source (28, 43, 44). These results showed that *Nha1* depletion resulted in increased defecation rate and excretory fluid loss as revealed by the production of more abundant, circular, larger and less concentrated excreta as compared to control-injected animals (Fig. *6D-H*). Indeed, the deposits produced by *Nha1* knockdown animals were often visibly associated with a striking increase in excess fluid as manifest by a ‘halo’ surrounding the excreta, which was never observed in control injected beetles (Fig. 6*I*). To test whether this increased excretory water loss was causally linked to defects in the reabsorptive capacity of the rectal complex, we adapted and optimized an *ex vivo* method based on isolating the entire alimentary canal under paraffin oil to quantify fluid reabsorption by the system (45); the preparations remained viable for several hours (*SI appendix*, Fig. S5). These experiments revealed that *Nha1* knockdown almost completely abolished fluid reabsorption by the rectal complex (Fig. 6*J*), thus demonstrating that this is the site responsible for the observed reduction in fluid retention. Taken together, our data suggest that loss of NHA1 function in leptophragmata cells dramatically impairs water reabsorption by the rectal complex, which affects systemic water balance and reduces the ability of adult beetles to survive desiccating conditions.

**Fig. 6.**
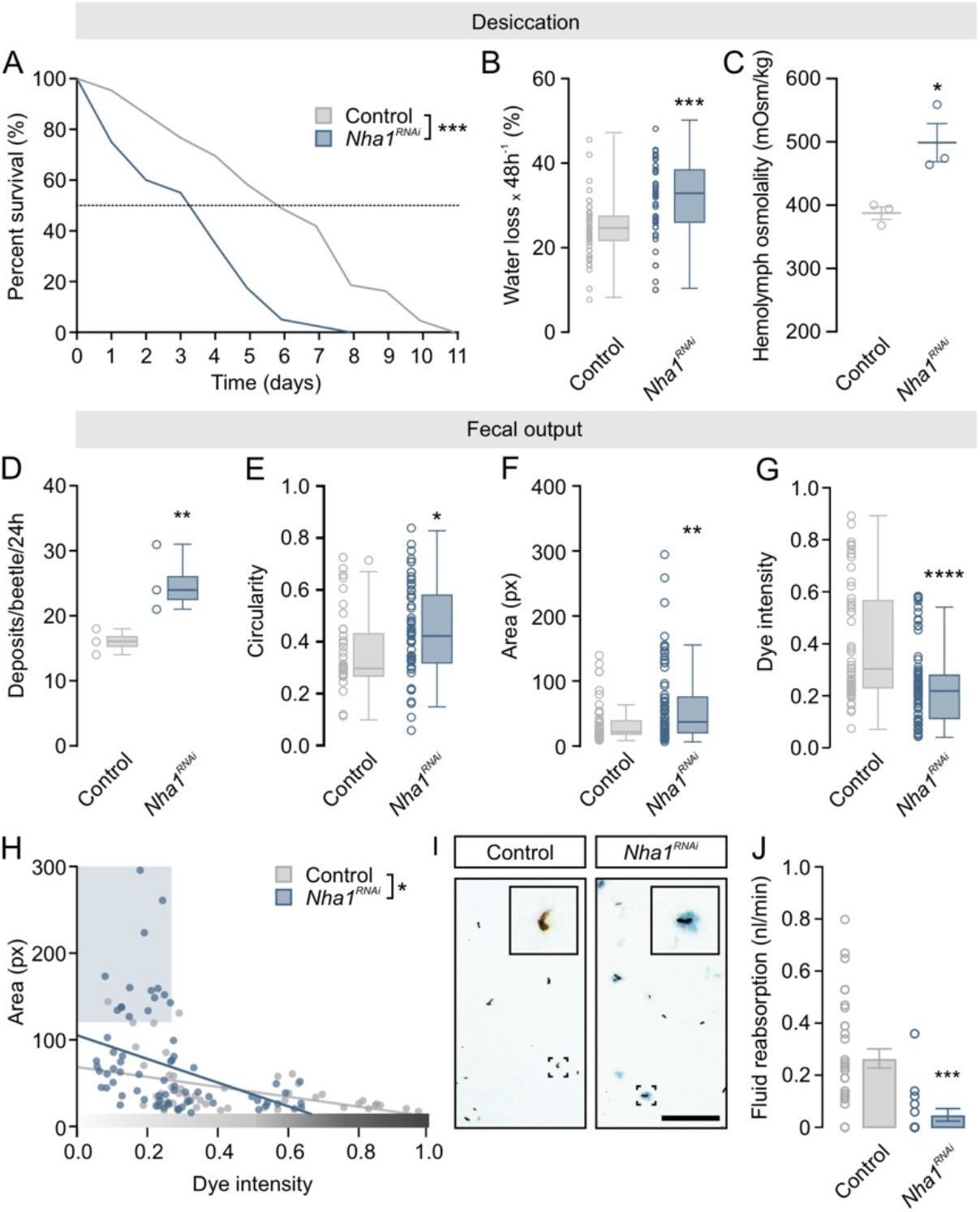
Systemic water balance depends on NHA1 activity. (*A*) Kaplan-Meyer survival function of adult control (dsRNA targeting *beta-lactamase, amp^R^*) and *Nha1* silenced animals. Adult specific knockdown of *Nha1* reduce organismal survival when exposed to low humidity conditions compared to control (RH 5%, log-rank test, *n* = 41-43). (*B*) Gravimetric analysis of control and *Nha1* silenced animals. Desiccation-induced water loss is significantly increased in *Nha1* knockdown animals relative to control (unpaired Student’s *t* test, *n* = 57, *** *P*<0.001). (*C*) Hemolymph osmotic pressure of control and *Nha1* depleted beetles. Hemolymph osmolality is significantly increased in *Nha1* knockdown animals relative to controls (unpaired Student’s *t* test, *n* = 3, *** *P*<0.05). (*D*) Defecation rate of *Nha1* depleted animals is significantly increased relative to controls (unpaired Student’s *t* test, *n* = 3 groups with 10 animals per group, ** *P*<0.05) and (*E*) the deposits are more circular (* *P*>0.05), (*F*) larger (** *P*<0.01), and (*G*) with a reduced dye intensity (**** *P*<0.001). (*D-G*) Statistical differences were tested using unpaired Student’s *t*-test. (*H*) The size of the deposits is inversely correlated with dye intensity in excreta from both *Nha1* silenced (blue trend line) and control animals (grey trend line) yet the slopes of the trend lines are significantly different (* *P*<0.05). Note the subpopulation of large, less intense deposits (blue shaded box), which is produced almost exclusively by *Nha1* knockdown beetles. (*I*) Representative images of excreta produced by control and *Nha1* knockdown animals. (Scale bar, 1 mm). (*J*) *Ex vivo* preparations of *Nha1* knockdown beetles show a significantly lower rectal complex-mediated fluid reabsorption rate relative to control (unpaired Student’s *t* test, *n* = 17-33, *** *P*<0.001).

### Tiptop-induced NHA1 expression underlies water-vapor absorption by the rectal complex

The powerful water-extracting properties of the rectal complex are not only related to the reabsorption of water from the feces, but can also be coupled to the remarkable ability to absorb water vapor directly from moist air (5–7, 11). Given that this mechanism depends critically on generating sufficiently low water activities in the rectal lumen to allow condensation of water from the atmosphere (6), we hypothesized that *Nha1*-knockdown would impair the ability to perform water vapor absorption. As predicted, we found that *Nha1*-deficient animals consistently lost body water when deprived of food at high humidity, whereas control animals retained, or even gradually increased, body water during identical exposures (Fig. 7*A-B*). These data confirm that *Tribolium* is able to absorb water at high relative humidities in a manner similar to that observed for *Tenebrio* (7, 11, 25)—albeit with a reduced capacity—and that this process depends critically on NHA1.

**Fig. 7.**
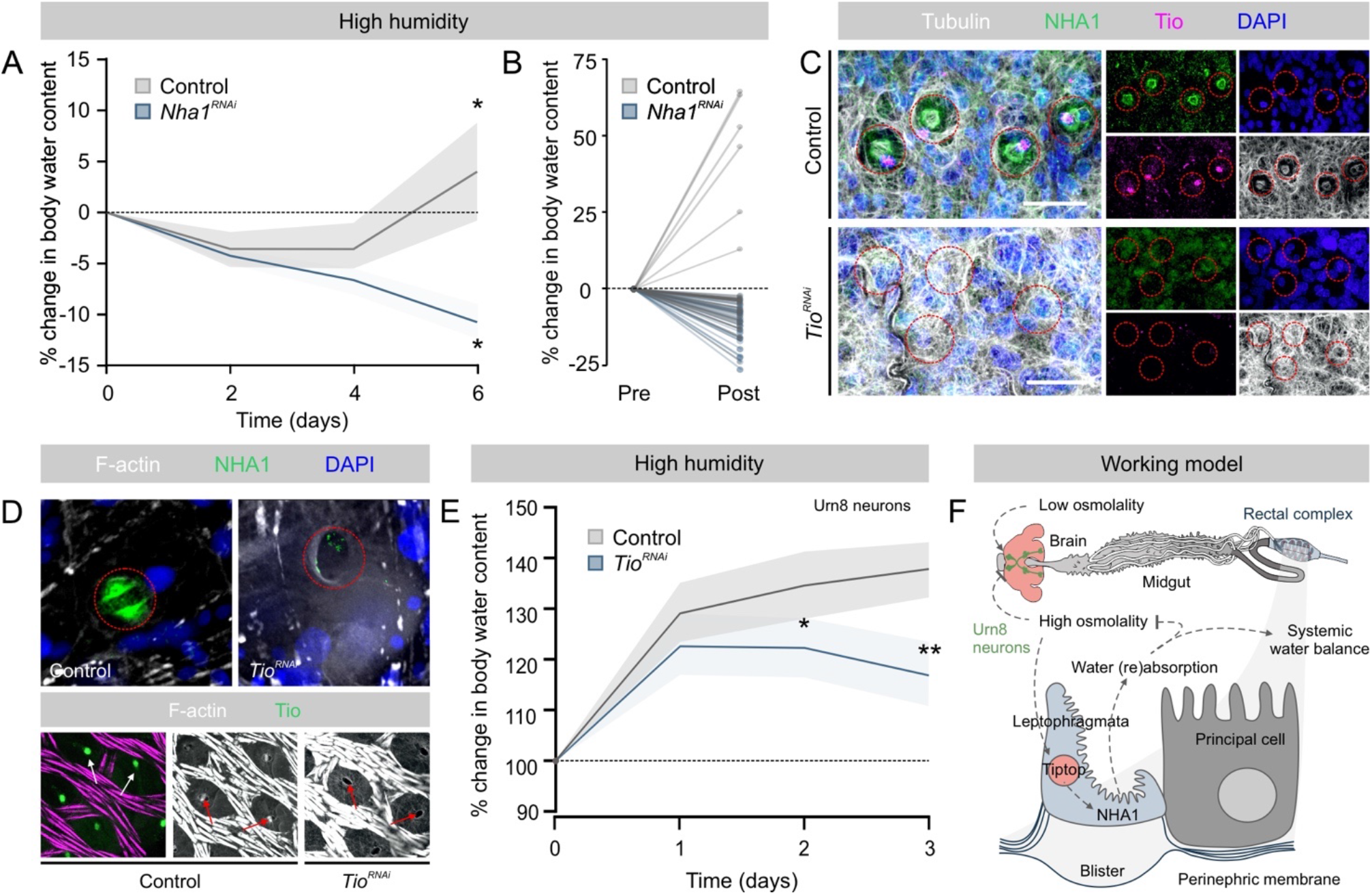
Tiptop controls NHA1 expression and leptophragmata differentiation. (*A*) Changes in body water in control and *Nha1* silenced animals exposed to RH 90% as a function of time. *Nha1* depleted animals consistently lose water over a six-day period, while controls retain, or even increase, organismal water levels relative to their initial water levels (paired Student’s *t* test, *n* = 40-49, * *P*<0.05). (*B*) Before-after plot of individual measurement graphed in (*A*) at day 0 (pre) and day 6 (post). (*C*) Immunolocalization of NHA1 and Tio in control and *Tio*-silenced animals. *Tio* knockdown results in almost complete loss of NHA1 expression as well as in morphological defects in the leptophragmata. (*D*) *Tio* depletion similarly abolishes NHA1 expression and causes defects in leptophragmata cytoarchitecture, which (*E*) significantly impairs water vapor absorption compared to controls in larvae of *Tenebrio* (unpaired Student’s *t* test, *n* = 3-7, * *P*<0.05; ** *P*<0.01). (*F*) Model for the systemic control of water balance in tenebrionid beetles.

We next explored the gene regulatory networks that govern *Nha1* expression and leptophragmata physiology. The finding that Tio—a transcription factor involved in secondary cell differentiation in many insects (28–30)—coexpressed with NHA1 in the leptophragmata of the PTs (Fig. 3*I*), suggests that Tio controls *Nha1* expression in leptophragmata. Accordingly, we selectively downregulated *Tio* expression during early development and subsequently probed for NHA1 expression in the rectal complex of adult beetles. The results revealed that *Tio* knockdown caused a complete loss of NHA1 expression, and further induced overt defects in the cytoarchitecture of the leptophragmata as evidenced by the failed formation of blister-like windows in the perinephric membrane (Fig. 7*C*). To test whether these defects affected whole-animal physiology, we silenced *Tio* expression in larvae of *Tenebrio* (the larger size makes them more amenable to gravimetric studies) and assessed their ability to absorb water vapor(6). As observed in *Tribolium*, knocking down *Tio* in *T. molitor* caused an almost complete loss of NHA1 expression as well as clear morphological changes to the leptophragmata of the rectal complex (Fig. 7*D*). Furthermore, silencing *Tio* expression consistently impaired their ability to absorb water as the body water contents of knockdown animals were significantly lower than mock-injected controls (Fig. *7E*). Taken together, our study identifies Tio as a key regulator of NHA1 expression and leptophragmata differentiation, which is essential to the function of the rectal complex and to maintenance of systemic water balance in *Tribolium* (Fig. 7*F*).

## Discussion

### The transport functions of the specialized leptophragmata underpin the water-conserving functions of the rectal complex

Classically, absorption of water by Tenebrionid beetles when exposed to subsaturated air was believed to occur across the cuticle, but it has since emerged that this process is entirely performed by the modified rectal complex. (6, 7, 12, 25). In this work we demonstrate that NHA1 in the specialized leptophragmata cells of the PTs is essential for the water-conserving functions of the rectal complex as well as the homeostatic control of water balance in *Tribolium* (Fig. 7*F*). How might we explain these physiological effects? Our combined analysis of the NHA1 protein suggest that, while it is capable of handling both Na^+^ and K^+^ movement, the main transport function of NHA1 is likely to mediate K^+^/H^+^exchange. If this transport modality is replicated in intact animals, these data strongly imply that NHA1 is functionally coupled to the active uptake of K^+^ by the complex, and thus to generating the osmotic forces necessary to facilitate water removal from the feces. Consistent with this idea *Nha1* depletion results in impaired fluid reabsorption by the rectal complex, increased excretion, and in reduced organismal water levels. Moreover, both NHA1 transcript and protein are upregulated in response to cues related to internal water stress. However, a difficulty presents itself when trying to reconcile these observed physiological effects with the fine structure of the rectal complex. Our work and previous studies (6, 7, 9) suggest that the main route of active KCl transport into the system is through the specialized leptophragmata, but these cells do not possess the anatomical hallmarks of active ion-transporting cells. The leptophragmata lack basal infoldings, have a reduced brush border, and contain few mitochondria (25, 46). It would therefore seem unlikely that these cells are solely responsible for mediating the high KCl concentrations measured in the PT lumen (6). By contrast, the larger PCs possess extensive basal infoldings, contain an extensive brush border full of mitochondria (7, 25) and are characterized by high expression of the plasma membrane V-ATPase (Fig. 3*G*). We therefore speculate that the H^+^ electrochemical gradient generated by the V-ATPase in the PCs could drive the secondary active transport of K^+^ via NHA1 in the leptophragmata, with Cl^-^ following passively (7) to produce the necessary osmotic gradients. Such a model is supported by the recent observation that luminal-directed K^+^ secretion is mediated by the SCs of the ‘free’ parts of the MTs in *Tribolium* (28), and is further analogous to that proposed for Na^+^ uptake by NHA2 in the SCs of MTs from the mosquito *Anopheles gambiae* (47). However, *Nha1* is not expressed in the ‘free’ parts of the tubule in *Tribolium* (BeetleAtlas.org), and Urn8R—a hormone receptor diagnostic of the SC identity—was not detectably expressed in the leptophragmata cells (28), implying that the PTs of the rectal complex operate via a two-cell type model that is distinct from that of the free tubules (28, 29). Mapping the exact routes and mechanisms with which water and ions flows through this multi-organ system remains an exciting prospect for the future. This could be addressed by single-cell RNAseq analysis of the rectal complex to obtain cellspecific insights into the molecular machinery that underlies the actions of the PT and rectal epithelia to help understand one of the most powerful water-conserving systems in biology.

### Exploring the transcriptional landscapes of *Tribolium*

Transcriptomic atlases are powerful tools in molecular genetics as they offer a detailed spatio-temporal view of gene expression that provides valuable clues to each gene’s physiological function (14, 48–50). Here, we introduce BeetleAtlas; a transcriptomic resource that provides a comprehensive view of the genetic signatures that underpin the functions of distinct tissues or life stages in the genetic model organism *Tribolium castaneum* (Fig. *1A-B*; *SI Appendix* Table S1). Besides being a useful addition to the rapidly expanding toolbox for *Tribolium* research, BeetleAtlas will broaden the focus of the beetle community. *Tribolium* is often adopted for studies in evolutionary development; but by cataloguing gene expression across different life stages, BeetleAtlas can help inform a much wider range of biological questions. This is significant because, although systemic RNAi is robust at all stages of development (51, 52), this technique precludes tissue-specific gene interference, and so the tissue(s) that contribute to a given phenotype remain unknown. Our online resource can thus provide a useful filter with which to identify the major tissue(s) in which candidate genes are most abundantly expressed and therefore most conveniently studied. Such an approach might be particularly useful for the *post hoc* analysis of data derived from large-scale functional screens (20, 53, 54). To further encourage this utility, BeetleAtlas is fully linked to existing databases (iBeetleBase (20) and FlyAtlas (14)) by a common vocabulary, enabling access to complementary information (e.g. RNAi phenotype and *Drosophila* ortholog relations) that will promote its use and integration in future studies.

In addition to allowing a simple gene-by-gene lookup, there is also scope for meta-analyses of the underlying transcriptomic datasets. For example, rather than adopting a candidate gene approach based on already known processes it is possible to use a hypothesis-free method by asking: “which genes are uniquely expressed in the adult brain, as compared to the larval brain?”. Such a methodology has the potential to generate unexpected research hypotheses independent of prior knowledge by allowing unbiased insights into the genes that underpin the functions of individual tissues or life stages (28, 55, 56). As well as offering a global analysis of gene expression there is further information to be extracted by querying the underlying relational database. The GAL4/UAS binary expression system is introduced in *Tribolium* (18), yet most GAL4 drivers may not be entirely specific to particular life stages of individual tissues. Genes with tissue-specific or ontogenetically restricted roles can be easily identified using BeetleAtlas (e.g. *SI Appendix*, Table 1), and used to make GAL4 driver lines with tissue- or stage-specific activities. Conversely, genes with ubiquitous expression or with persistent roles throughout development can also be found. In sum, we predict that this online resource will have a major impact on the insect functional genomics community by providing an extensive catalogue of gene expression across different tissues and life stages in *Tribolium*, which will promote both an evolutionary as well as an ontogenetic and tissue-centered view of gene function.

## Materials and Methods

### Animal Husbandry

*Tribolium castaneum* (San Bernardino strain) stocks were maintained on organic wholemeal wheat flour supplemented with 5% (w/w) yeast powder (*Tribolium* medium) at 30°C at a constant 50% relative humidity (RH) and 12:12 light-dark cycles as in (57). *Tenebrio molitor* was cultured on organic bran supplemented with occasional potato slices under identical environmental conditions.

### Tissue Dissection and RNA Extraction

Tissues were dissected from non-sedated 6th instar larvae or 1-week-old mature adults under a freshly prepared mixture of Schneider’s medium (Invitrogen, CA, US) and *Tribolium* saline (1:1, v/v). The *Tribolium* saline contained: NaCl 90 mmol l^-1^, KCl 50 mmol l^-1^, MgCl_2_ 5 mmol l^-1^, CaCl_2_ 2 mmol l^-1^, NaHCO_3_ 6 mmol l^-1^, NaH_2_PO_4_ 6 mmol l^-1^, Glucose 50 mmol l^-1^ and the pH was adjusted to 7.0. Dissected tissues were then transferred to 500 μl QIAzol (Qiagen, Hilden, DE) and stored at –80°C until sufficient tissue had been collected to allow extraction of a minimum of 100 ng RNA in total. Next, the samples were thawed and physically disrupted using a beadmill (1 min max speed) using a TissueLyser LT (Qiagen, Hilden, DE) and then extracted with phenol-chloroform including an extra chloroform step and several RNA washing steps. The RNA was then finally purified using a Qiagen RNeasy Plus mini kit according to the manufacturer’s instructions. The optional DNase step, the optional drying of the column, and back-elution were all included. For each sample, the concentration of RNA was determined using a NanoDrop 1000 Spectrophotometer (ThermoFisher, MA, USA), and the quality of the RNA was determined using an Experion Pro260 (Bio-Rad, CA, USA) with Experion RNA HighSens Analysis Kit (BioRad, CA, USA), according to the manufacturer’s instructions. Each tissue sample was prepared in biological triplicates.

### RNA-seq Analyses and Database construction

Total RNA libraries were prepared for each sample according to a low-input protocol by BGI Genomics (Shenzhen, Guangdong, China), and sequenced on a BGISEQ-500 using paired-end chemistry (100 nt reads) with a sequencing depth of 6 Gb per sample (i.e. approaching 40x of the ~150 Mb *Tribolium castaneum* genome). The resulting fastQ.gz files were processed through the Tuxedo pipeline (58) using version 5.2 of the *T. castaneum* reference genome assembly (59), and the output used to populate a MySQL relational database, entitled TriboliumDB. The database also contains gene data from the Tcas 5.3 reference genome, gene ontology information from the Gene Ontology Consortium (www.geneontology.org), and *Drosophila melanogaster* symbol and name information from FlyBase (flybase.org).

The database, TriboliumDB, underlies a web application, BeetleAtlas, publicly available at www.BeetleAtlas.org. The web application employs a Java servlet to generate web pages and communicate with the TriboliumDB database, and separate smaller servlets for subsidiary functions. It contains a documentation (‘Docs’) section with full details and version dates. As a web application, BeetleAtlas thus allows non-technical users to make a wide range of prepared queries to the underlying relational TriboliumDB. This database is freely available for download, so that more sophisticated custom queries can be made, if required, by those with access to basic bioinformatics expertise. For this study, the BeetleAtlas web application was interrogated for gene orthologs of known ion channels and transporters (‘Gene’ lookup function) as well as for genes enriched in the rectal complex relative to the whole-animal signal (‘Tissue’ enrichment function), with all candidate genes prioritized according to enrichment.

### Scanning electron microscopy (SEM)

SEM analysis of the rectal complex was performed according to a modified protocol described in (55). In brief, rectal complexes were dissected under Schneider’s medium and briefly exposed to a AgNO3 solution (30 sec) as described in (27) before being fixed in 2.5% glutaraldehyde in 0.1M cacodylate buffer (pH 7.4) for 90 min as in (43). The tissue was then rinsed repeatedly in ddH2O before being dehydrated through a graded ethanol series, and desiccated using an Autosamdri-815 critical point dryer (Tousimis Research Corporation, Maryland, USA). The rectal complexes were then transferred to aluminum stubs, fractioned and coated with platinum (70 s B12 nm thickness) in a JEOL JFC-2300HR high-resolution fine coater (Jeol, Tokyo, Japan) and examined with a Zeiss Sigma variable pressure scanning electron microscope (Carl Zeiss, Oberkochen, Germany) using secondary electron (SE) and back-scatter electron (BSE) detection methods to sequentially visualize both the topology and element weight distribution (atomic number, *Z*) of the samples.

### Antibody Generation and immunolocalization of target proteins

To generate specific antibodies against proteins of interest, we analyzed the amino acid (aa) sequence of the proteins to identify the best immunizing peptide region according to a previously described method (60). For NHA1, this analysis resulted in the selection of a peptide corresponding to aa 547–562 (SMSTTVSQKDSPKGE) in the C-terminal region of the full-length parent protein, which was then submitted for a custom immunization protocol carried out by Genosphere Biotechnologies (Paris, France). Additionally, aa 491-496 (PAATLAEFYPRDSRH) of the VHA55 peptide was also selected for preparation of polyclonal antisera. Epitope specificity of the different antisera was established by comparing wild-type and RNAi animals by immunostaining.

Immunohistochemistry on paraffin sections was carried out as in (61). Briefly, rectal complexes were dissected and fixed in 4% paraformaldehyde in PBS for 30 min before being dehydrated in a graded series of ethanol (70%, 90%, 99%) followed by incubation in xylene for 20 min. Next the tissues were incubated in paraffin for 30 min (3x exchanges with fresh paraffin) and embedded in paraffin wax in a suitable mold and left to cool for 2 days. Then, semithin sections were cut on a Leica ultramicrotome EM UC6 (Leica Microsystems, Wetzlar, Germany) with glass knives and mounted on objective slides. Slides were then dewaxed in Histo-Clear (Scientific Laboratory Supplies, US) for 15 min, hydrated in a decreasing series of ethanol (99%, 90%, 70%), and finally washed in ddH_2_O for 30 sec. Slides were finally stored in PBS and immunostained using out anti-VHA55 antibody as described below.

Immunocytochemistry (ICC) was performed as in (55). In brief, rectal complexes were dissected and fixed as described above. Tissues were then washed four-six times in PBST (PBS + 0.1% Triton X-100), blocked with PBST containing 2 % normal goat serum (blockPBST; Sigma-Aldrich, MO, USA) for 1 h, and incubated in primary antibodies. Primary antibodies used were polyclonal rabbit anti-NHA1 (1:500), polyclonal mouse α-VHA55 (1:500) and polyclonal rat α-Tio (62) (1:500). The subcellular location of the endogenous proteins were visualized by applying Alexa Fluor 488/647 anti-rabbit, anti-mouse or anti-rat secondary antibodies (1:500; Sigma Aldrich, MO, USA) in combination with DAPI (1:1000) and Rhodamine-conjugated Phalloidin (1:500; Sigma Aldrich, MO, USA) in blockPBST overnight at 4°C. Following several washes, first in PBST and then in PBS, the different tissues were mounted on poly-L-lysine coated 35mm glass bottom dishes (MatTek Corporation, MA, USA) in Vectashield (Vector Laboratories Inc., CA, USA) and imaged on an inverted Zeiss LSM900 confocal microscope equipped with airy scan 2 technology (Zeiss, Oberkochen, Germany). Where necessary, immunofluorescence was quantified using the FIJI software package from images acquired using identical microscope settings as described in (43).

### Computational Modeling and NHA1 Structure-Function Predictions

The 3-dimensional tertiary structure of NHA1 was predicted by using I-TASSER (35) and a best-fit model was selected based on its confidence score (C-score). Further, Ramachandran Plot Assessment (RAMPAGE) was utilized to calculate the torsional angles and side chain conformations of all amino acid residues contained in the NHA1 protein sequence to perform structural refinement of the model. Finally, we performed dynamic stability estimations of NHA1 in the presence of either Na^+^ or K^+^ ions, by submitting the protein for 10 ns molecular dynamics (MD) simulation using AMBER18 (63) on the Computerome 2.0 high-performance computing cluster. System preparation including minimization and equilibrium processe were performed as previously described (64). The CPPTRAJ module incorporated in AMBER suite was used for trajectory analysis according to their root mean square deviation (RMSD), radius of gyration (RoG), and hydrogen bond calculations. During the entire simulation period, plots presented the comparative stability of modelled NHA1 in the presence of Na^+^ or K^+^relative to each other.

### Molecular Cloning of *Nha1*

cDNA of *Nha1* was synthesized from total RNA extracted from adult *T. castaneum* rectal complexes using the High-Capacity cDNA Reverse Transcription Kit with RNase Inhibitor (ThermoFisher, MA, USA), and the coding region of the gene was amplified using Q5^®^ Hot Start High-Fidelity 2X Master Mix (New England Biolabs, MA, USA) using *Nha1*-specific primers (Supplemental Table 2). The PCR products were subsequently cloned into pGEMHE *Xenopus laevis* oocyte expression vector using In-Fusion^®^ HD cloning kit (TaKaRa Bio Inc, Kusatsu, JP) and the final sequence validated (Eurofin, Luxemburg, LU).

### pH Measurements in *Xenopus laevis* Oocytes

cRNA was generated from *Nha1* in PGEMHE using mMessage mMachine (Ambion, Austin, TX, USA). *X. laevis* oocytes (Ecocyte Bioscience, Dortmund, Germany) were injected with 50 nl cRNA solution containing 25 ng *Nha1* cRNA per oocyte using a microinjector (Nanoject, Drummond Broomall). The oocytes were incubated 4-7 days hours at 19°C in Kulori’s solution (90 mM NaCl, 4 mM KCl, 1 mM MgCl_2_, 1 mM CaCl_2_, 5 mM HEPES, pH 7.4) before making pH recordings. pH-electrodes were from borosilicate glass capillaries with filament (120F-3, WPI) using a vertical puller (Narishige Scientific Instrument Lab). The resistance of the pipettes was 3-4 MOhm when filled with 3 M KCl and submerged in Kulori’s solution. The pipettes were baked for 2 hr at 220 °C in a metal box. Dimethyldichlorosilane (Silanization Solution I, Sigma Aldrich) was added to the box though the hole in the lid and the pipettes were silanized at 220 °C for 2-4 hr. The tip of the silanized pipettes was filled with a H^+^-selective ionophore cocktail (hydrogen ionophore I cocktail A, Sigma-Aldrich) by dipping the tip of the pipette into the solution. The electrodes were back-filled with a solution containing 40 mM KH_2_PO_4_, 23 mM NaOH and 150 mM NaCl, pH 7.5. Next, a pH electrode was mounted on the head stage of an EPC7 amplifier, and calibrated using 100 mM KCl with 10 mM MES/TRIS pH 5.5, pH 6.6 and pH 7.5 before each recording and after each recording. The range was typically 55-60 mV/pH. If the voltage had shifted more than 5 mV, the recording was discharged. *X. laevis* oocytes were placed in a recording chamber and superfused with a standard Kulori’s solution and impaled with the pH electrode. After 10 min the standard Kulori’s solution was replaced by 1) a sodium free Kulori’s solution where NaCl was substituted by choline-Cl (0Na^+^) for 10 min then by Kulori’s solution, pH 8.5 (pH 8.5) for 10 min. 2) Kulori’s solution was replaced by 15 mM NaCl and 90 mM KCl, 1 mM MgCl_2_, 1 mM CaCl_2_ og 5 mM HEPES, pH 7.4 (90K^+^) for approx. 5 min and the superfused by the same solution with KCl substituted with cholin-Cl (0K^+^). The effects of the different solutions on the membrane potential was tested and two electrode voltage clamp were done using an Oocyte Clamp OC-725B (Warner Instruments). The pipettes were filled with 3 M KCl solution and the reference electrodes were connected to the bath via agar bridges with 3 M KCl. The signal in the voltage recording electrode was subtracted from the signal in the pH electrode to calculate the pH potential based on the calibration curves.

### Gene expression analysis

Validation of RNAi-mediated gene knockdown and environmentally induced changes in gene expression was assessed by quantitative Real-Time PCR (qPCR). Total RNA extraction was carried out 3 days after dsRNA injection and cDNA synthesis were carried out as described above. Next, qPCR was performed using the QuantiTect SYBR Green PCR Kit (Fisher Scientific, NH, USA) in combination with a Stratagene Mx3005P qPCR system (Agilent Technologies, CA, USA). The effect of *Urn8* depletion and DH37 hormone stimulation on *Nha1* expression was also assessed. This was done by injecting (25 nl volume) either PBS or PBS containing DH37 peptide corresponding to a final peptide concentration of approx. 10^-7^M into adult animals with samples collected at 3 hours and 6 hours after treatment. Expression levels were normalized against the housekeeping gene *rp49*. All primers used are listed in Supplemental Table 2.

### Production of dsRNA and RNAi-mediated Knockdown

To silence target gene expression by RNAi, transcript sequences covering app. 200-500 bp were selected. Total RNA was then extracted from rectal complexes (showing highest enrichment of *Nha1*) and cDNA synthesis was carried out as described above. Using the cDNA as template, fragments were amplified by PCR using gene-specific primers that were tagged with T7 promoter sequences at both the 3’ and 5’ ends (see Supplemental Table 2). These gene-specific fragments were then cloned into the pUC19 vector individually and subsequently verified by sequencing (Eurofins, Luxembourg, L). Using the cloned vector as template, bidirectional *in vitro* transcription was carried out using the MEGAscript T7 transcription kit (ThermoFishcer, MA, USA), and the quality of the resulting dsRNA was checked by gel electrophoresis and quantified using NanoDrop. The concentration was adjusted to 2 μg/μl using injection buffer (1.4 mM NaCl, 0.07 mM Na_2_HPO_4_, 0.03 mM KH_2_PO_4_, 4 mM KCl), and a total of 500 nl dsRNA solution was injected into age-matched adults using a Nanoject II injector (Drummond Scientific, PA, USA). Animals were allowed to recover for 2 days after injection before being used for experimentation.

### Environmental stress exposure

In control (fed) conditions, beetles were housed individually in a 96-well plate with standard wholemeal flour containing 5% yeast. For drinking-only (water) treatments, animals were kept in 96-well plates with a small block of 1% agar with 0.05% bromophenol blue (BPB). Drinking was verified by the presence of blue deposits. For desiccation treatments, animals were kept in 96-well plates with a piece of filter paper without any nutritional and water sources at 30°C, with individual plates kept at 5%, 50% or 90% RH.

### Desiccation tolerance

Animals were kept on *Tribolium* medium for 3 days after dsRNA injection. Healthy animals were then transferred to a 96-well plate in a container filled with silica gel beads (Sigma-Aldrich, MO, USA) to produce a low-humidity environment (approx. RH 5%, measured by a custom-build hygrometer). The number of dead animals (not responding to tactile stimuli) were then counted every 8 h for 7 days. Data were expressed as percent survival over time.

### Hemolymph collection and quantification

Hemolymph was collected according to the protocol described in (28, 43, 65) from animals exposed to the different environmental stress exposures as described above. In brief, animals were washed and subsequently dried on tissue paper for 2 hours to remove moisture. Then, beetles were anesthetized by CO_2_ and their cuticle pierced between the pronotum and elytron before being transferred to an ice-cold 0.5-ml tube with a small hole in the bottom in groups of 10. This tube was then placed in a larger 1.5-ml collecting tube, which was centrifuged at 12,000 x *g* for 15 min at 4°C. Hemolymph from separate tubes were combined into each collecting tubes (containing 500 μl paraffin oil to prevent oxygen-induced melanization) from each environmental condition. Following sample collection, each sample was diluted to a final volume of 50 μl with ddH2O and the osmotic pressure of each sample was measured in triplicates on VAPRO Vapor Pressure Osmometer Model 5600 (Wescor Inc., UT, USA) with each measurement corrected according to the dilution factor of the sample.

### Quantification of water content

To measure changes in total water content, individual beetles were transferred to a small plastic container and then measured on a Sartorius SE2 ultra micro balance (=W_T_; Sartorius, Göttingen, DE; 0.1 μg readability). The animals were then housed under low humidity conditions as described above, and after 48 h the beetles were reweighted (= W_48_). To measure the corresponding dry weight of the animals, they were kept at −20°C over night and then placed in a 65°C incubator for at least 2 days before being weighed a final time (=W_dry_). The percent water loss of total body water for each animal was calculated as (W_T_ – W_T48_)/(W_T_ – W_dry_) × 100%, with *N*=25 animals weighed for each experimental group.

### Defecation Behavior

To assess the effects of manipulating *Nha1* expression on whole-animal excretory behavior *in vivo*, dsRNA-injected animals were starved for 2 days followed by refeeding a standard *Tribolium* medium supplemented with 0.05% (w/w) Bromophenol blue (BPB) sodium salt (Sigma-Aldrich, MO, USA) overnight as in (28). This special medium was created by mixing the standard *Tribolium* medium with BPB and a small amount of water hereby creating a uniform paste, which was left to dry at room temperature overnight. The dried BPB-labelled *Tribolium* medium was then ground to a fine powder creating a consistency identical to that of the standard medium. Beetles were then placed in individual wells of a 96-well plate fitted with a small piece of filter paper and the number of BPB-labelled deposits produced by each animal over a 4 h period was quantified.

### *Ex-vivo* fluid reabsorption assay

The water reabsorption rate from the CNC *ex vivo* was assessed using a modified protocol (45). The control and *Nha1-KD* animals were dissected carefully, keeping the head, gut, tubules and CNC intact under Schneider’s medium. MTs were then broken from the entry point (common trunk) to the CNC. The dissected animals were carefully transferred to another dish containing paraffin oil (molecular grade) with a wax layer at the bottom, and gently stretched (to keep intact the fore-, mid-, hindgut, MTs, and CNC) and pinned at the head and anal cuticle. Two drops (5μl and 0.3μl) of premixed 3x *Tribolium* saline + Schneider’s solution (1:1) supplemented with 100 μmol l^-1^ of amaranth (Sigma-Aldrich, St Louis, MO, USA) were dropped under paraffin oil and their circumference measured by formula using the eye-piece graticule (10 mm = 200 parts) (Leica Microsystems, Germany). A 5μl drop was pulled around the midgut portion and MTs and a 0.3μl drop onto the CNC with the help of a capillary pull glass rod, avoiding contact with the head and cuticle. MTs were gently drawn out from the saline drop and then the open ends of MTs were wrapped around the pin with the help of fine forceps. This was left for two hours—the maximum time the system remained stable. The change in circumference of the initial and final drop was measured as V = (π x d^3^)/6, and the rate of absorbance was calculated by the following formula: *J_fluid_* =Δv/Δt where *J_fluid_* is the fluid reabsorption rate (nl min^-1^), Δv is the change in volume (nl), and Δt is the duration of the experiment (min).

### Water vapor absorption assay

The ability to extract water vapor directly from the atmosphere was quantified gravimetrically as described in (5). In brief, animals were desiccated for 2 days before being weighed and individually housed in a 96-well plate without food or water. The plate was then placed in a high-humidity chamber (RH >95%) in a temperature-controlled incubator and the weight of each animal recorded once a day for 3-6 consecutive days (depending on species) after which the animals were sacrificed, dried for 2 days at 60°C, and the reweighed to calculate the changes in body water over the time, as described above.

### Statistics

The statistical analyses were performed using the data analysis software GraphPad Prism 9 (CA, USA). The normal (Gaussian) distribution of data were tested using the D-Agostino-Pearsen omnibus normality test. Data were plotted as mean ± SEM, Tukey’s box-and-whisker plots or as raincloud plots, as indicated in each figure legend. Statistical differences between one control group and another group (unpaired samples) or between the same groups at different time points (paired samples) were compared using the two-tailed Student *t*-test, whereas differences between one control group and several other groups were pairwise compared by one-way ANOVA followed by Dunnett’s multiple comparisons tests taking P=0.05 (two-tailed) as the critical value. P-values are indicated as: * P < 0.05, ** P < 0.01, *** P < 0.001, **** P < 0.0001.

## Acknowledgements

We are grateful for Gregor Bucher for providing *Tribolium* stocks and embryo samples. This work was supported by the Villum Foundation (grant no 15365), by the Danish Council for Independent Research (grant no. 9064-00009B) as well as by Ragna Rask-Nielsens Foundation (KH0622) and research infrastructure grant by Carlsberg Foundation (grant no. CF19-0353) to K.V.H. Further support was given by The Carnegie Trust (grant no. 70425) and Leverhulme Trust to B.D. (grant no. RPG-2019-167). The authors declare no conflicts of interest.

## Figures and Tables

**Fig.S1.**
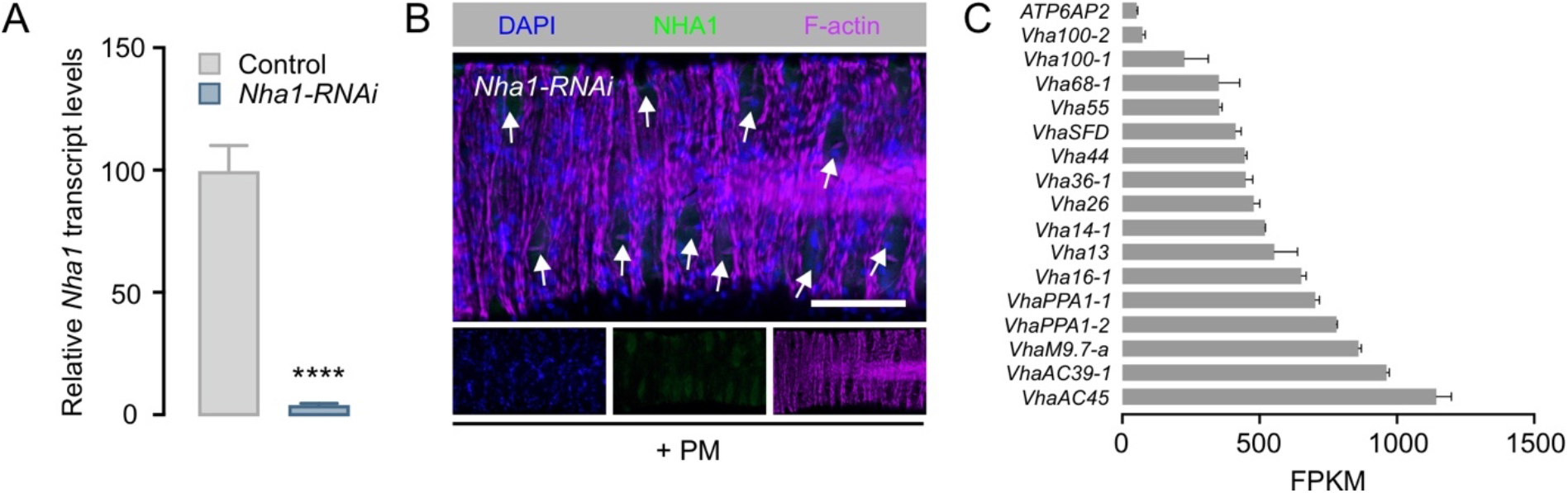
Validation of *Nha1* knockdown efficacy. (*A*) *Nha1* transcript levels in the rectal complex (*n* = 5) from animals injected with dsRNA targeting the *Nha1* gene (*Nha1-RNAi*) show a significant knockdown of Nha1 expression relative to mock inject controls (unpaired Student’s *t* test, **** P<0.0001). (*B*) Rectal complexes dissected from *Nha1*-depleted animals show almost complete depletion of anti-Nha1 immunoreactivity (arrows). (Scale bar, 100 μm). PM, perinephric membrane. (*C*) Transcript levels of V-ATPase subunit genes in the rectal complex obtained from BeetleAtlas.org.

**Fig.S2.**
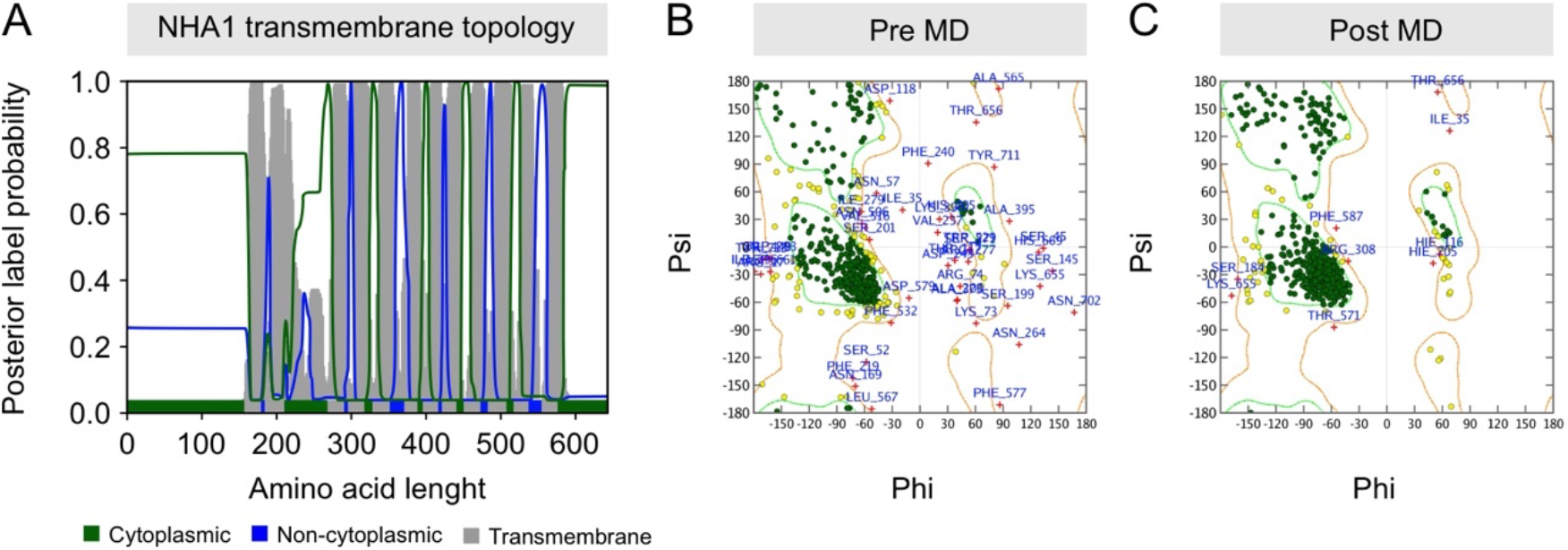
Structural analyses of NHA1 protein. (*A*) Transmembrane topology of NHA1 revealing an 11-transmembrane domain with a tertiary structure, characteristic of other CPA2 family members. (*B*) Ramachandran plots of pre-molecular dynamics and (*C*) post-molecular dynamics (MD) simulation analyses of NHA1. The post-MD modelled structure shows an improved refinement and accuracy relative to the pre-MD structure, thus validating the use of the post-MD structure for stability analyses.

**Fig.S3.**
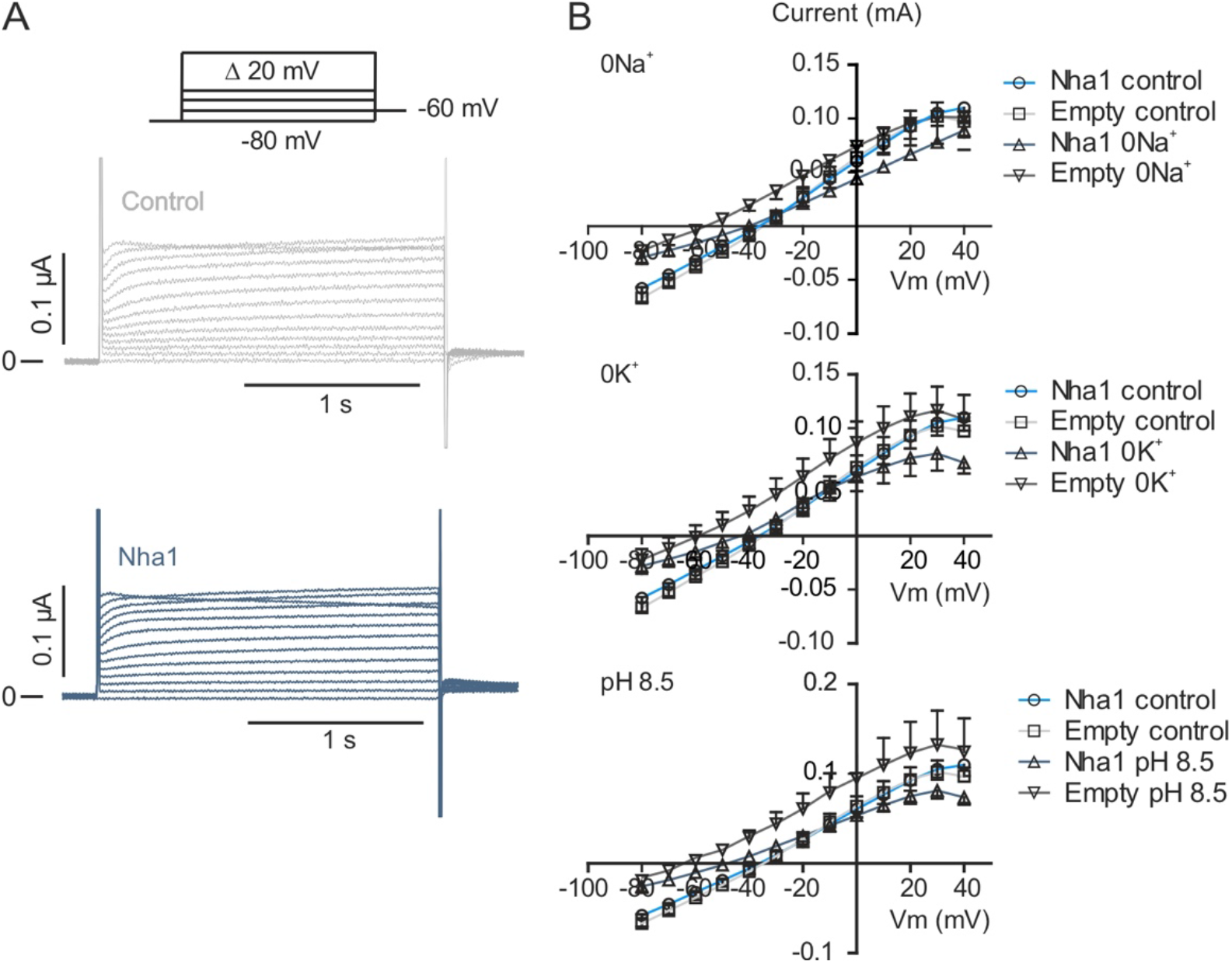
NHA1is electroneutral and does not exhibit voltage-dependence. *Nha1* was expressed in *Xenopus laevis* oocytes and compared to uninjected oocytes (Empty control). Currents activated by the depicted voltage-clamp protocol in Kulori’s solution (control), in 0 mM K^+^, 0 mM Na^+^ and in pH 8.5. (*A*) Voltage-clamp protocol and representative currents recorded from empty controls and *Nha1* injected oocytes. (*B*) Mean currents plotted as a function of voltage, n=6-8.

**Fig.S4.**
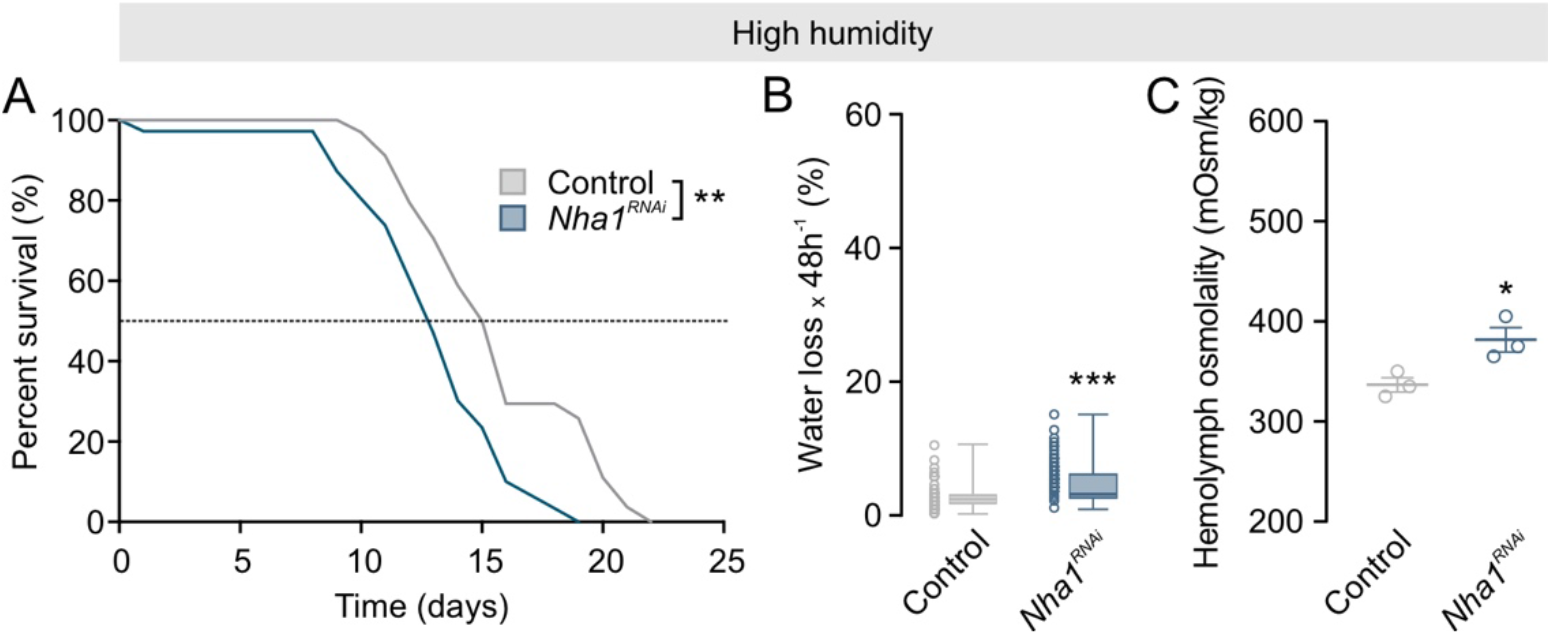
*Nha1* depletion affects systemic water balance during high humidity exposure. (*A*) Kaplan-Meyer survival function of control and *Nha1* silenced animals. Knockdown of *Nha1* reduces organismal survival when exposed to higher humidity compared to the control (RH 90%, log-rank test, *n* = 39-44). (*B*) Gravimetric analysis of control and *Nha1*-silenced animals. Water loss is significantly increased in *Nha1* knockdown animals relative to control at high humidity (unpaired Student’s *t* test, *n* = 57, *** *P*<0.001). (*C*) Hemolymph osmotic pressure of control and *Nha1*-depleted beetles. Hemolymph osmolality is significantly increased in *Nha1* knockdown animals relative to controls at high humidity (unpaired Student’s *t* test, *n* = 3, *** *P*<0.05).

**Fig.S5.**
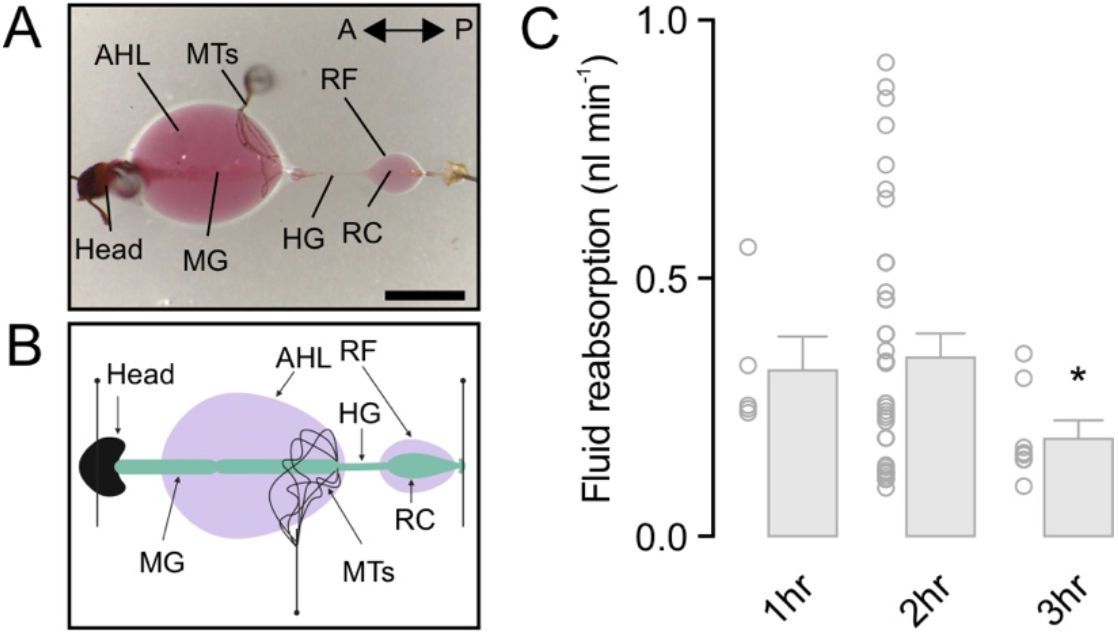
Modified *ex vivo* fluid reabsorption assay. (*A*) Light-micrograph and (*B*) schematic representation of the *ex vivo* fluid reabsorption assay of the rectal complex. (*C*) Rate of reabsorption of fluid by the rectal complex with time. The rate of reabsorption of fluid is significantly reduced after 3 hr compared to the rate measured after 1hr (One-way ANOVA, *n* = 17-33, * *P*<0.05). AHL, artificial hemolymph solution; HG, hindgut; MTs, Malpighian tubules; MG, midgut; RC, rectal complex; RF, rectal reabsorbed fluid. Scale bar 0.6 mm.

**Table S1.**
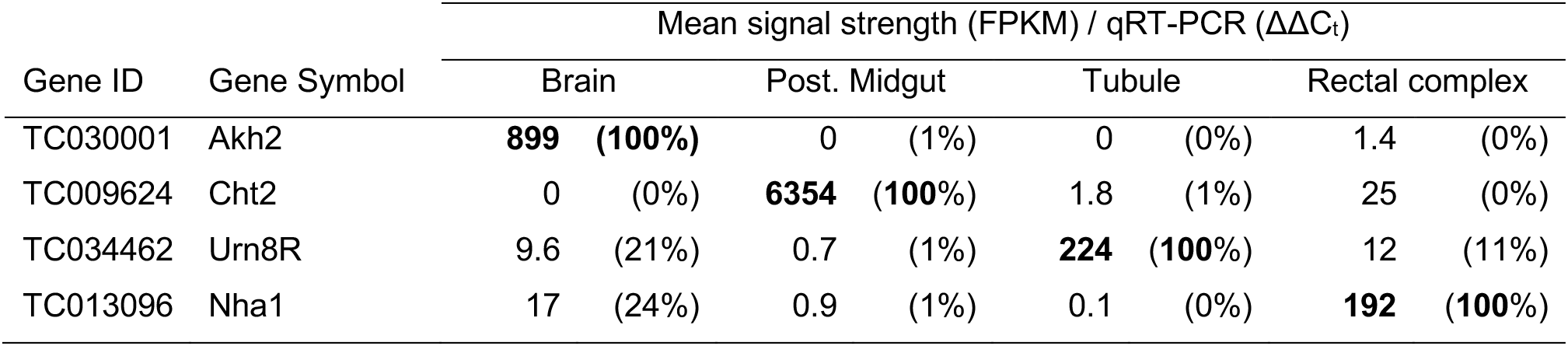
Examples of genes showing highly tissue-specific expression patterns according to BeetleAtlas, thus validating the specificity and quality of the RNAseq data. These data are further independently validated by qRT-PCR. Boldface indicates the maximum signal for each gene.

**Table S2.**
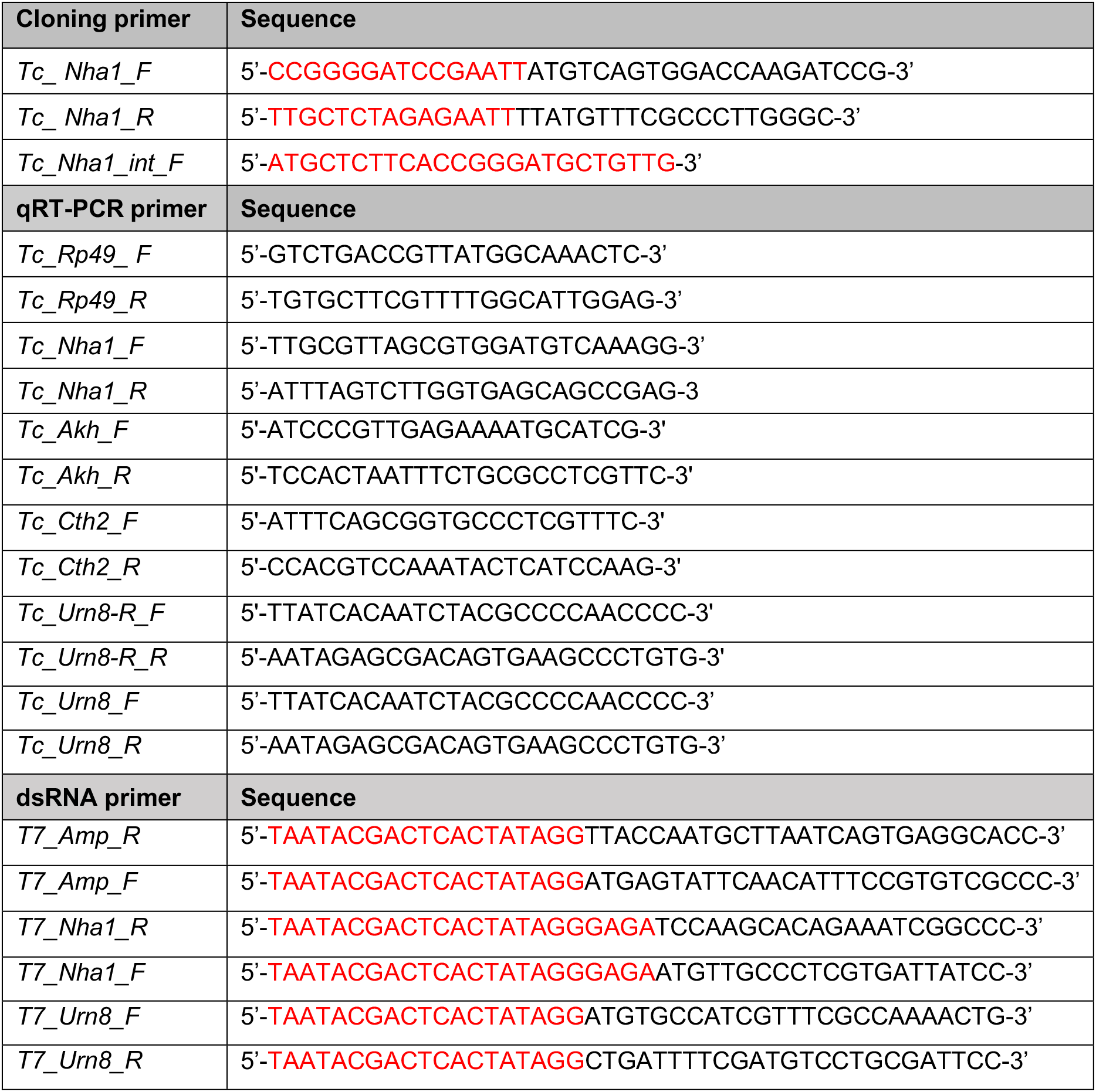
Primer sequences used for In-Fusion cloning, RT-qPCR and dsRNA synthesis. Sequences marked in red correspond to the vector sequence for In-Fusion cloning primers and to the T7 promoter sequence for dsRNA primers.

